# Ambroxol and Ciprofloxacin Show Activity Against SARS-CoV2 in Vero E6 Cells at Clinically-Relevant Concentrations

**DOI:** 10.1101/2020.08.11.245100

**Authors:** Steven B Bradfute, Chunyan Ye, Elizabeth C Clarke, Suresh Kumar, Graham S Timmins, Vojo Deretic

## Abstract

We studied the activity of a range of weakly basic and moderately lipophilic drugs against SARS CoV2 in Vero E6 cells, using Vero E6 survival, qPCR of viral genome and plaque forming assays. No clear relationship between their weakly basic and hydrophobic nature upon their activity was observed. However, the approved drugs ambroxol and ciprofloxacin showed potent activity at concentrations that are clinically relevant and within their known safety profiles, and so may provide potentially useful agents for preclinical and clinical studies in COVID-19.

## Introduction

The novel severe acute respiratory syndrome coronavirus 2 like virus, (SARS-CoV2), ^1,2^ was identified as the causative agent of the current pandemic of viral pneumonia, causing a disease termed COVID-19 (<u>cor</u>ona<u>vi</u>rus <u>d</u>isease-20<u>19</u>) by the World Health Organization, with efficient human to human transmission, and significant morbidity and mortality. The world is scrambling to develop countermeasures, including therapeutics aiming to lessen disease severity and to develop with prophylaxes including vaccines.

Recently, FDA approved drugs, chloroquine (CQ) and hydroxychloroquine (HCQ) and azithromycin (AZT) have shown therapeutic activity against COVID-19 in initial clinical trials,^3^ and also in *in vitro* cellular systems, although subsequent clinical evidence does not suggest these are effective. All these agents are known to inhibit autophagy in humans at relevant doses,^4–7^ and so the field is actively looking to repurpose and develop other modulators of autophagy that may potentially show anti SARS-CoV-2 activity. Another possible mechanism we have proposed is related to their weakly basic and hydrophobic nature, whereby they may modify pH in acidified intracellular sites such as endosomes and the trans-Golgi network,^**8–10**^ although this has yet to be validated in the context of SARS-CoV-2 infected cells. Here, we present data upon the activity of these drugs, and also other weak bases we have studied, ciprofloxacin (CPX),^10^ ambroxol (AMB), and its prodrug bromhexine (BHX) against SARS-CoV2 in Vero E6 cells. Unlike the inhibition of autophagy caused by CQ, HCQ, AZT and CPX^11^, both BHX^12^ and AMB^13^ induce execution of the complete autophagy pathway from initiation to maturation at concentrations that are clinically relevant.

## Materials and Methods

### Biosafety

All procedures involving live virus were performed in a BSL3 facility with the approval of the institutional biosafety committee.

### Cell death inhibition assay

Vero E6 cells were treated with the indicated compounds for 1 hour prior to infection with 0.05 MOI of SARS-CoV-2 Isolate USA-WA1/2020 (deposited by the Centers for Disease Control and Prevention and obtained through BEI Resources, NIAID, NIH, NR-52281). Forty-eight hours later, cell viability was assessed by XTT Cell Viability (ThermoFisher) using the manufacturer’s protocol. Data are means +/− standard error (SE) of 3 replicates. At least one additional replicate experiment was performed for each compound.

### Quantitative PCR assay

Supernatants from cells infected as above were harvested and RNA was isolated using the Qiagen Viral RNA Mini kit following the manufacturer’s protocol. Primers and probes as designed by the Centers for Disease Control and Prevention were obtained from Integrated DNA Technologies 2019-nCoV RUO Kit (Cat # 10006713). qPCR was performed using the TaqMan™ Fast Virus 1-Step Master Mix (Applied Biosystems). Assays were conducted on a QuantStudio5 Q-PCR machine (Applied Biosystems), and reported as cycle threshold values (Ct). Data are means +/− standard error (SE) of 3 replicates. At least one additional replicate experiment was performed for each compound (except CPX/AMB mixtures), and representative data shown.

### Plaque Forming Unit Assay

Supernatants from infected cells as above were added to fresh Vero E6 cells and incubated for 2 hours at 37°C and aspirated. Cells were overlaid with 1 mL of a mix of 2% agarose and 2X minimal essential medium with 2.5% FCS and incubated at 37°C for 2 days, followed by fixation with 4% formaldehyde. After overlay removal, cells were stained with 0.5% crystal violet, washed, and dried. Plaques were counted for determination of viral titer.

### Statistical Analysis

Analyses were performed with GraphPad (GraphPad Prism 7.0) and Sigma Stat (Sigma Stat 3.0).

## Results

The first assay used was based upon survival of Vero E6 cells after infection with SARS-CoV2, so that antiviral effects will manifest themselves as increases in cell survival, commonly referred as protection against cytopathic effect. For the autophagy-inhibiting weak base agents, robust and dose-dependent increases in survival were observed for both CQ and CPX, while AZT had no effect (Figure 1). For the autophagy inducing weak bases a dose dependent increase in survival was observed for AMB but not for BHX (Figs. 1 and 2). The second assay used was quantitative PCR (qPCR) of SARS-CoV2 genome to determine the effect of replication, so that antiviral effects will manifest themselves as increases in cycle threshold value (Ct). For the autophagy inhibiting weak bases, AZT at 10 μM had no effect on Ct, 10 μM CQ showed a moderate increase in Ct, while CPX caused a marked and dose-dependent increase in Ct (Fig 3). For the autophagy inducing weak bases, AMB showed a dose-dependent increase in Ct, with highest activity at 100 μM. (Fig. 3). When AMB (10 or 100 μM) and CPX (10 μM) were combined, the effects appeared antagonistic. Finally, the third assay used was a plaque formation assay to confirm that the qPCR results are indicative of true infective viral units (Fig 4). CPX showed a marked and dose dependent decrease in viral titer (Fig. 4), the highest concentrations of AMB and CQ showed modest reductions, and AZT showed no effect.

**Figure 1.**
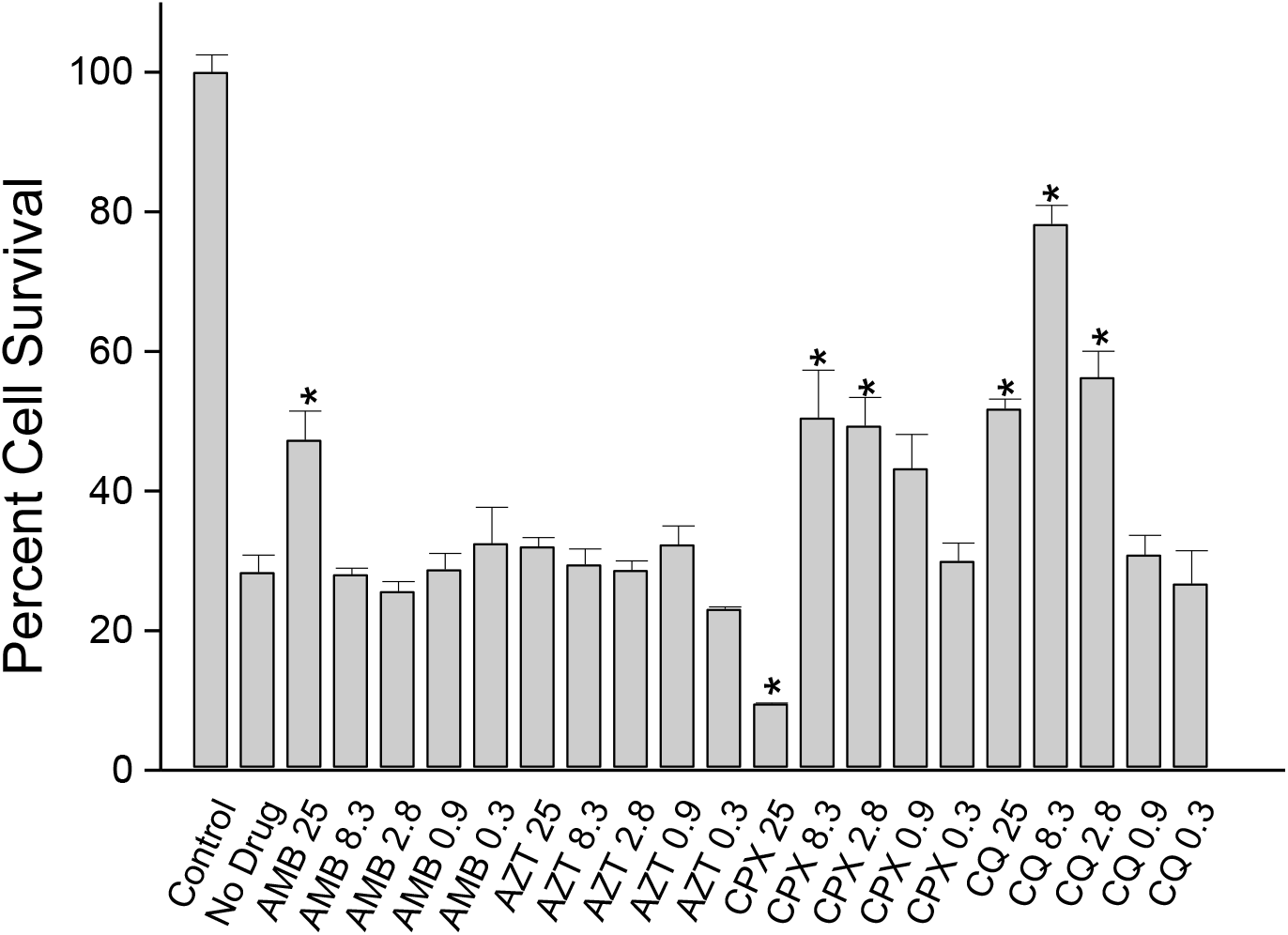
The effect of compounds on Vero E6 cell survival 48 hours after treatment with SARS CoV2. Control cells were uninfected, and represented 100% survival. Data are means of n= 3 +/− SE from a representative of two separate experiments. Cells were treated with compounds at the micromolar concentrations shown. * Denotes significantly different (p < 0.05) from no drug sample by ANOVA.

**Figure 2.**
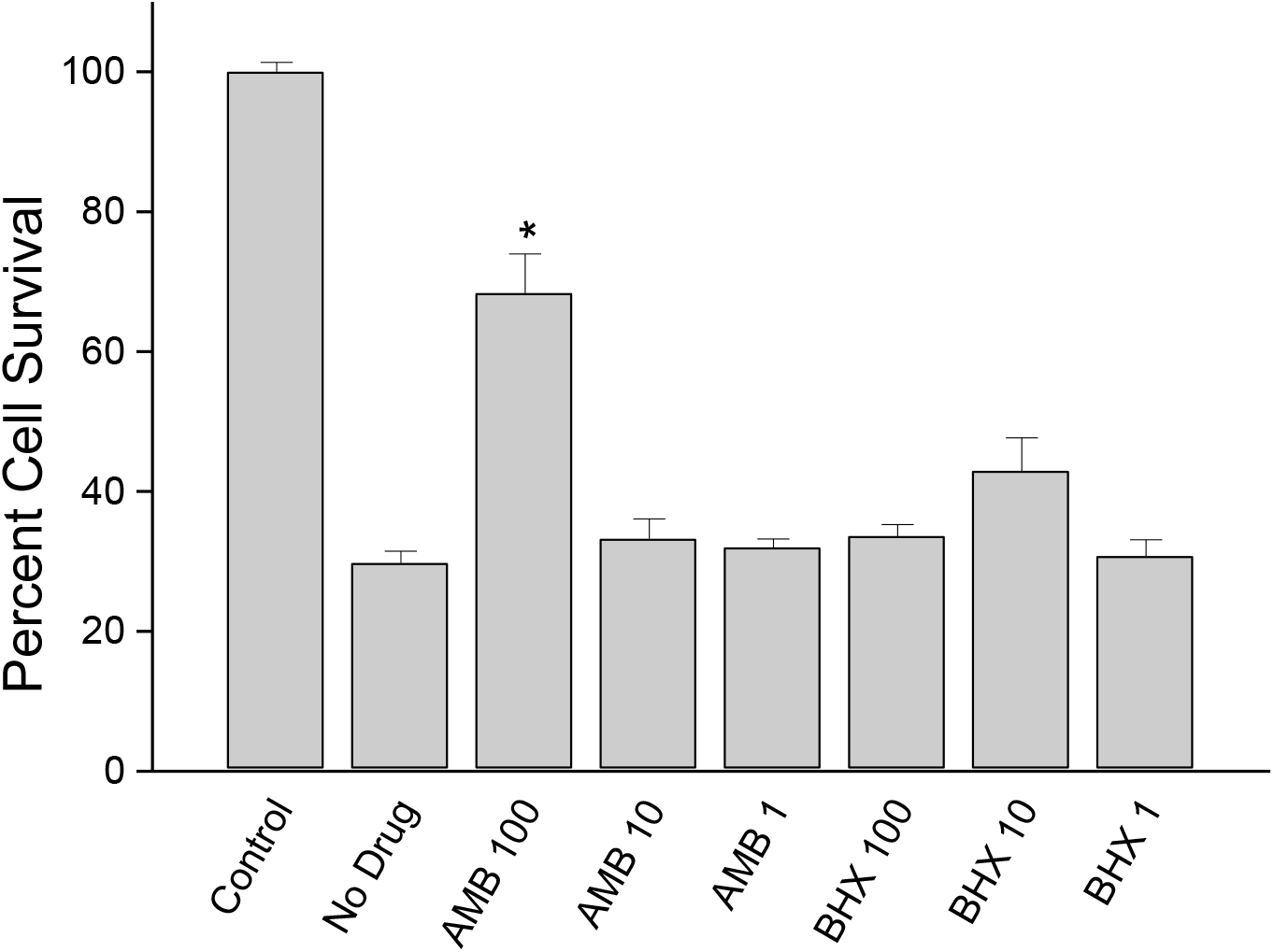
The effect of ambroxol (AMB) and bromhexine (BHX) on Vero E6 cell survival 48 hours after treatment with SARS CoV2. Control cells were uninfected, and represented 100% survival. Data are means of n= 3 +/− SE from a representative of two separate experiments. Cells were treated with compounds at the micromolar concentrations shown. * Denotes significantly different (p < 0.05) from no drug sample by ANOVA.

**Figure 3.**
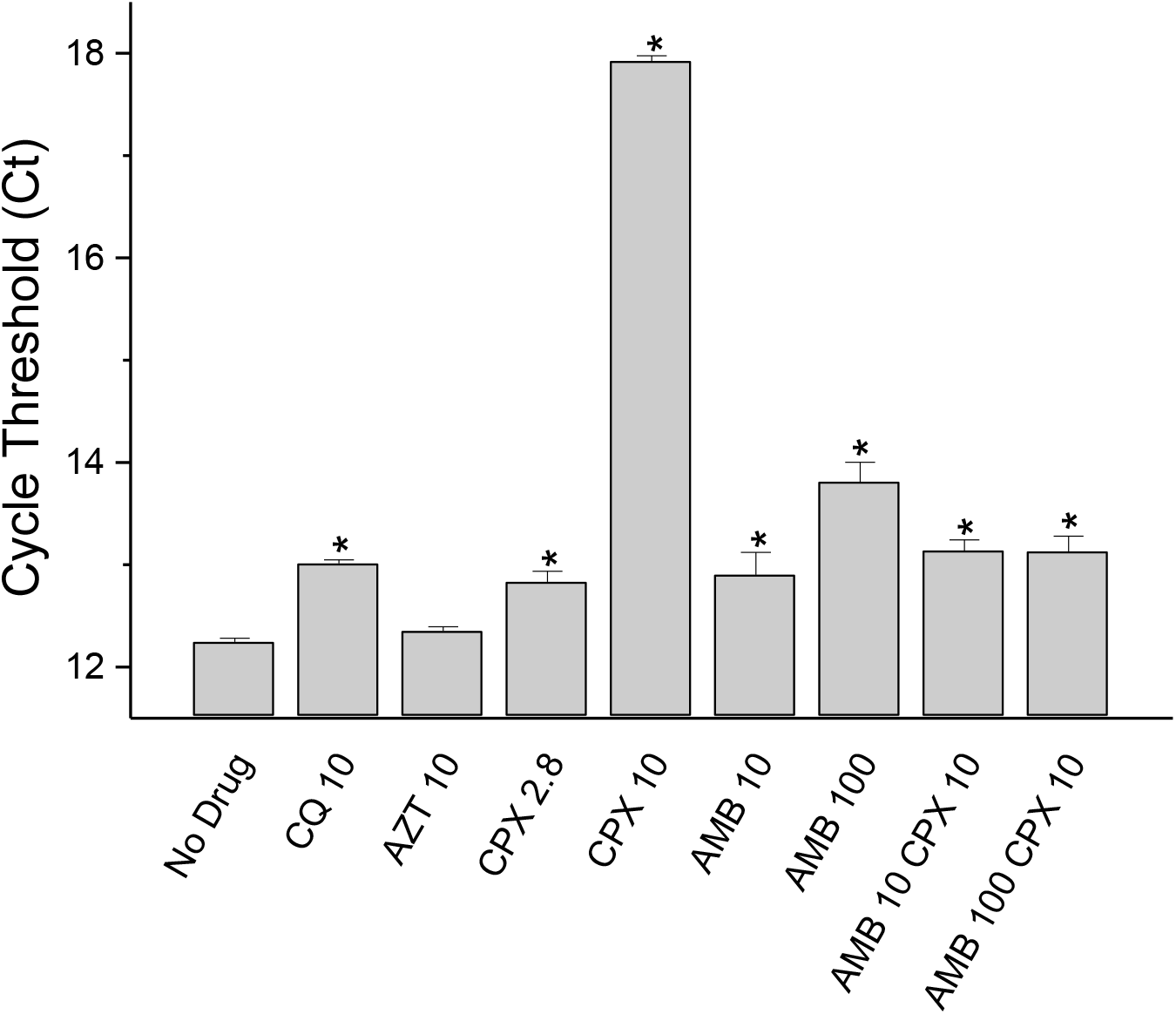
The effect of compounds on the cycle thresholds (Ct) for qPCR of viral RNA in supernatents from Vero E6 after treatment with SARS CoV2. Control cells were uninfected, and represented 100% survival. Data are means of n= 3 +/− SE from a representative of two separate experiments (AMB/CPX combination was from a single experiment). Cells were treated with compounds at the micromolar concentrations shown. * Denotes significantly different (p < 0.05) from no drug sample by ANOVA.

**Figure 4.**
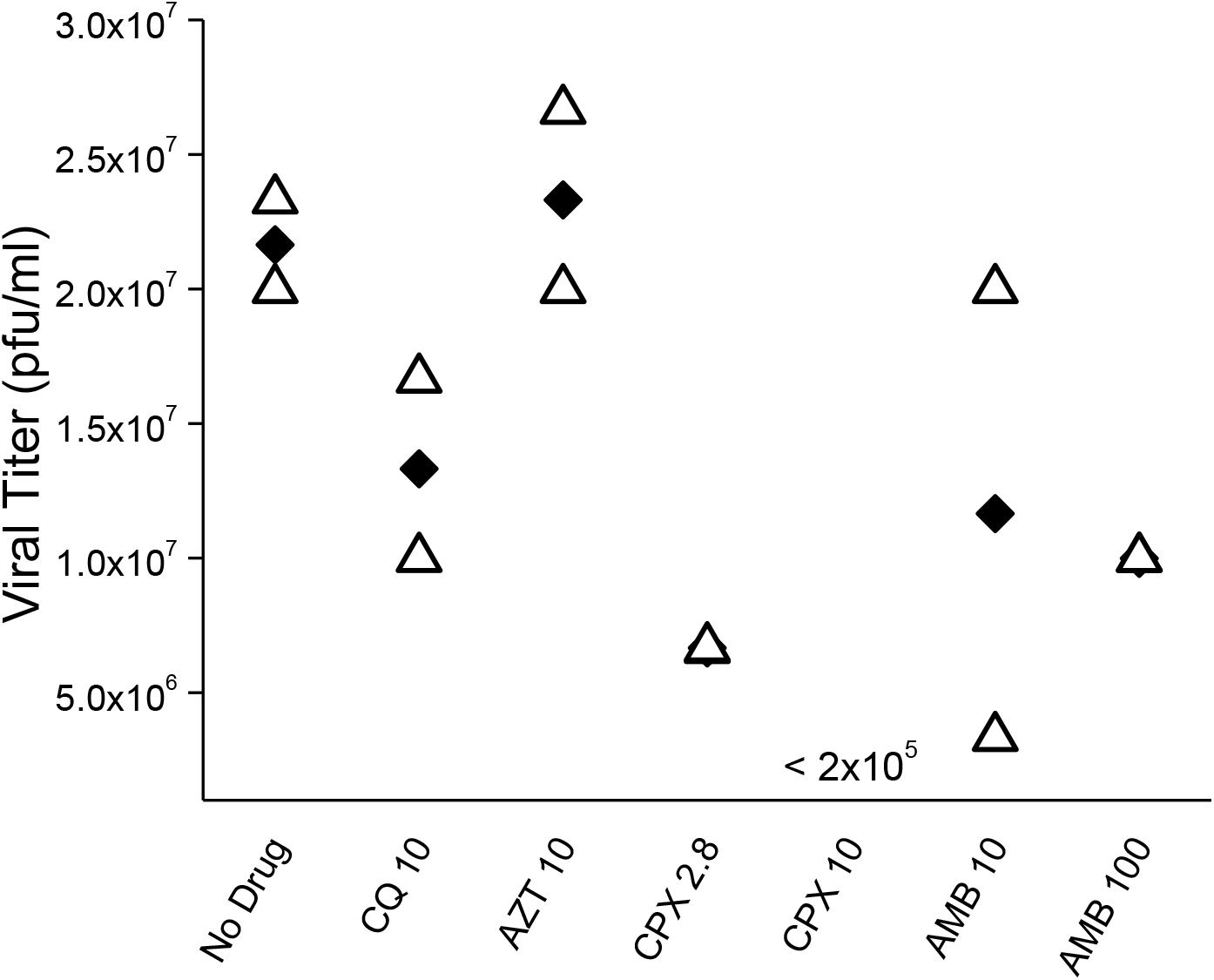
The effect of compounds on plaque forming units per milliliter (pfu/ml) in supernatants from Vero E6 after 48 treatment with SARS CoV2. Data are shown as average (solid diamond) and range (triangle), n=2 from a single experiment. Cells were treated with compounds at the micromolar concentrations shown. The data for 10 μM CPX were below the level of detection in this assay, 10^5^ pfu/ml.

## Discussion

Our initial hypothesis was based around the neutralization of organelle acidification,^**8–10**^ enabled by the weakly basic and lipophilic properties of the compounds used (Table 1). Despite broad similarities in their physical-chemical properties, there was no broad effect of all compounds on Vero E6 survival or viral replication, with AMB, CPX and CQ showing activity while AZT and BHX did not. Nor was there any clear correlation of activity depending upon autophagy status: of the inhibitors, CPX and CQ showed activity, while the inducer AMB also showed effects. Therefore, while their weakly basic and lipophilic nature may enable all these drugs to access and neutralize acidified organelles, this does not explain the anti-SARS-CoV2 activity of some of these, nor the absence of activity of others, in Vero E6 cells. Thus, additional mechanisms of action, including those that are directly anti-viral or affecting entry or other aspects of SARS-CoV-2 replication, transcription, and propagation, are postulated here.

**Table 1.** Selected properties of weak base drugs used. Not performed, n.d.

| Drug | logP | Amine pKa | Autophagic effect | Effect on Vero E6 Survival | Effect of SARS-CoV2 replication |
| --- | --- | --- | --- | --- | --- |
| AZT | 4.0 | 8.2, 8.6 | Inhibits | - | - |
| CPX | 1.3 | 8.7 | Inhibits | +++ | +++ |
| CQ | 4.6 | 10.1 | Inhibits | +++ | ++ |
| AMB | 2.6 | 9.0 | Induces | ++ | ++ |
| BHX | 4.3 | 9.3 | Induces | - | *n.d.* |

The activities of AMB and CPX were achieved at concentrations that are relevant to the lung tissue levels reached at known safe therapeutic doses. AMB is highly safe in both adult and pediatric use.^14,15^ It has also been shown safe in long term use at high doses in Parkinson’s disease (1260mg q24h)^16^ and Gaucher disease (27mg/kg q24h)^17^ and doses of this magnitude are known to lead to lung levels approaching 100 μM, the most effective concentration in this work, because of the potent lung tropism of this compound in humans.^18^ Furthermore, high dose AMB (15-20 mg/kg/day) has been shown both safe and also to reduce inflammatory cytokines in acute respiratory distress syndrome (ARDS),^19^ and so potential trials in COVID-19 of high dose AMB may be warranted.

Mean CPX lung levels after a 500 mg dose were have been observed as being above the most effective concentration here, 10 μM, for up to 4 hours after dosage,^20^ again demonstrating that the concentrations used here are clinically relevant. However, although a more potent agent in our hands than AMB, CPX has black box warnings for several adverse events. However, if its’ clinical activity is as superior as shown here in cells, its risk/benefit ratio may still be favorable, as both risk and benefit are higher. Retrospective analysis in COVID-19 cases treated with CPX (in secondary lung infection for example) should be conducted for determination of potential benefit. Since an inhaled form of CPX has been developed (Linhaliq reached Phase 3), this may lower systemic exposure while maintaining efficacy. The effects of combined AMB and CPZ on viral replication appear antagonistic, although additional studies are required.

AMB appears to have a separate and distinct activity from the TMPRSS2-dependent activity of BHX, and so the combination of these two drugs (high dose AMB and regular dose BHX), although counter-intuitive as BHX is the prodrug of AMB, may prove useful. This is especially so, as AMB is known to be safe at much higher doses than BHX, with these high doses also showing effects in ARDS that would be highly desirable in COVID-19 patients; increasing oxygenation (PaO_2_/FiO_2_ ratio, PO_2_ and SaO_2_) while also reducing inflammatory markers (TNF-α and IL-6).^19^

Mechanistically, BHX is a potent inhibitor of the protease TMPRSS2, ^21^ whose activity is necessary for SARS-CoV2 entry into many cell types,^22^ including respiratory epithelia, and so its lack of effects here might be considered puzzling. However, recent work shows that Vero E6 cells do not express TMPRSS2, and instead viral entry is achieved through an alternative and CQ-inhibited pathway .^23^ Thus, BHX may show benefits in cells where SARS-CoV-2 entry depends on TMPRSS2. However, in our hands, AMB appears to have a separate and distinct antiviral activity from that of BHX, with BHX being hypothesized^24^ and trialed^25^ as a repurposed treatment for COVID-19. Although the mechanisms of action of AMB and CPX against SARS-CoV2 in Vero E6 cells remain to be elucidated, there are some possibilities. In our previous studies, AMB was more potent than BHX in inducing autophagy^12,13^ and so it is possible that this reflects its higher activity here, or other target(s) may be important. Fluoroquinolones such as CPX act against susceptible bacteria by preferentially binding to single strand microbial nucleic acid,^26^ and this then forming an inhibitory ternary complex with either DNA gyrase or topoisomerase; the activity of CPX observed here may result from a similar inhibitory ternary complex with viral RNA and viral proteins such as helicase or RNA polymerase.

## Acknowledgements

VD was supported by NIH grants R37AI042999 and R01AI111935 and center grant P20GM121176.

## Study Strengths

The activities of AMB and CPX were confirmed in three independent assays and were achieved at clinically-relevant concentrations.

## Study Weaknesses

Only one cell line, Vero E6, was used. Vero E6 cells do not express TMPRSS2, a potential target of BHX.

